# Unique T cell signatures associated with reduced *Chlamydia trachomatis* reinfection in a highly exposed cohort

**DOI:** 10.1101/2023.08.02.551709

**Authors:** Kacy S Yount, Chi-Jane Chen, Avinash Kollipara, Chuwen Liu, Neha V Mokashi, Xiaojing Zheng, C Bruce Bagwell, Taylor B Poston, Harold C Wiesenfeld, Sharon L Hillier, Catherine M O’Connell, Natalie Stanley, Toni Darville

## Abstract

*Chlamydia trachomatis* (CT) is the most common bacterial sexually transmitted infection globally. Understanding natural immunity to CT will inform vaccine design. This study aimed to profile immune cells and associated functional features in CT-infected women, and determine immune profiles associated with reduced risk of ascended endometrial CT infection and CT reinfection. PBMCs from CT-exposed women were profiled by mass cytometry and random forest models identified key features that distinguish outcomes. CT+ participants exhibited higher frequencies of CD4+ Th2, Th17, and Th17 DN CD4 T effector memory (TEM) cells than uninfected participants with decreased expression of T cell activation and differentiation markers. No significant differences were detected between women with or without endometrial CT infection. Participants who remained follow-up negative (FU-) showed higher frequencies of CD4 T central memory (TCM) Th1, Th17, Th1/17, and Th17 DN but reduced CD4 TEM Th2 cells than FU+ participants. Expression of markers associated with central memory and Th17 lineage were increased on T cell subsets among FU- participants. These data indicate that peripheral T cells exhibit distinct features associated with resistance to CT reinfection. The highly plastic Th17 lineage appears to contribute to protection. Addressing these immune nuances could promote efficacy of CT vaccines.

**GRAPHICAL ABSTRACT:** 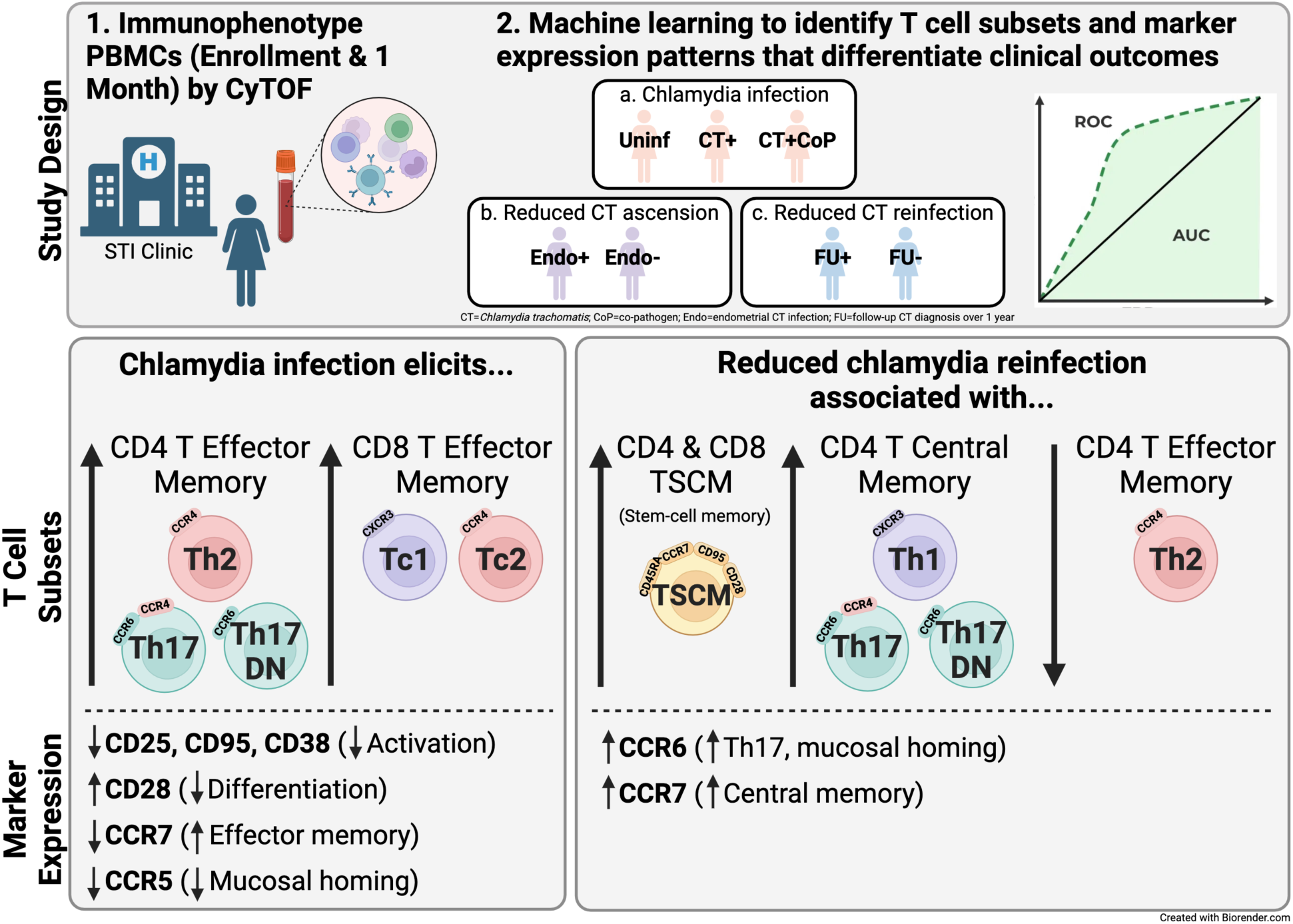

## INTRODUCTION

*Chlamydia trachomatis* (CT) is the most common bacterial sexually transmitted infection (STI) globally (1). In women, CT infects the cervix and can ascend to the uterus and oviducts to cause pelvic inflammatory disease (PID) that can result in pelvic pain, infertility, and ectopic pregnancy (2). Although antibiotic treatment is effective, 80% of infections are asymptomatic and may go undetected (3), contributing to the risk for chronic sequelae (4). Although repeated CT infections are common, they can elicit partially protective immunity. Women who spontaneously cleared infection without treatment were less likely to become reinfected (5) and persons with confirmed prior infection had lower chlamydial loads (6). Determining the aspects of natural immunity that drive protection against CT will advance understanding of chlamydial immunity and aid design of an effective vaccine.

Mouse models have elucidated protective mechanisms of different arms of the immune response. The importance of CD4 T cell mediated immunity is revealed from studies in severe combined immune deficient (SCID), RAG-1-/-, TCRα-/-, CD4-/-, and MHCII-/- mice, which display delayed or failed resolution of *Chlamydia* infection, while MHCI-deficient (B2M-/-), B cell-deficient (μMT), and CD8-/- mice resolve infection with comparable kinetics to wild-type (7–11). Th1 cells secrete IFNγ and TNFα that promote recruitment and activation of macrophages. IFNγ also induces expression of indoleamine 2,3-dioxygenase (IDO) in epithelial cells, which catabolizes tryptophan, an essential nutrient for chlamydia growth (12). IFNγ-knockout, antibody depletion, and adoptive transfer studies in mice confirmed that IFNγ-producing CD4 T cells (Th1) are key mediators of protection against *Chlamydia* (13–15), although this may be independent of T-bet, the defining transcription factor expressed by Th1 cells (16). Th2 cells produce IL-4, IL-5, and IL- 13 that drive antibody responses, but a *Ct-*specific Th2 clone did not protect against murine genital infection and displayed reduced trafficking to the genital mucosa, whereas a polyfunctional Th1 clone mediated chlamydial clearance (17, 18). Classical Th17 cells produce IL-17, recruit neutrophils, promote secretory IgA, and are instrumental in protection against many mucosal pathogens, but can be pathogenic. Mice lacking T-bet had Th17-skewed responses and similar clearance of *C. muridarum* infection to wild-type (16) while those lacking RORγt (Th17-deficient) exhibited delayed clearance (19), highlighting a potential protective role for Th17. However, we found that IFNγ-deficient mice had chronic pathology related to a compensatory increase in Th17 cells and downstream induction of neutrophils (20), highlighting the importance for a balance in Th17 responses.

The T Cell Response Against Chlamydia (TRAC) cohort is a unique resource containing clinical samples and information from women highly exposed to CT and other STIs that was established to improve understanding of human natural immunity to CT (21). In this cohort, we observed that peripheral blood CD4 T cells producing CT-specific IFNγ were associated with protection from reinfection (22), consistent with findings of others (23), and we additionally determined that CD8 T cells producing CT*-*specific IFNγ were associated with protection from endometrial infection (22). Further, we identified a single nucleotide polymorphism in the gene for CD151, a tetrospanin molecule important for immune cell adhesion and migration and T cell proliferation, as a candidate genetic marker for reduced susceptibility to endometrial infection (24). In contrast, we determined that anti-CT antibodies were insufficient to protect participants from endometrial or recurrent infection (25, 26). These data indicate that CD4 T cells, and to a lesser extent CD8 T cells, contribute to protection against CT reinfection or ascension.

The objective of this study was to immunophenotype peripheral blood from TRAC participants to identify specific T cell subsets and surface markers associated with T cell function that associated with resistance to CT reinfection and/or endometrial infection. Therefore, a CyTOF panel containing 33 markers was designed to deeply profile T-helper cell subsets such as T follicular helper (Tfh; CXCR5+) cells, regulatory T (Treg; CTLA4+ CD25+ CD127lo) cells, naïve T cells (TN; CD45RA+ CCR7+), central memory T (TCM; CD45RA- CCR7+) cells, effector memory T (TEM; CD45RA- CCR7-) cells, terminally differentiated T effector (TEMRA; CD45RA+ CCR7-) cells, and stem-cell like memory T (TSCM; CD45RA+ CCR7+ CD28+ CD95+) cells. TEM and TCM T-helper cell subsets were defined by surface expression of chemokine receptors CCR6, CXCR3, and CCR4 (27) because multiple studies have shown that CCR6-CXCR3+CCR4- (Th1), CCR6-CXCR3-CCR4+ (Th2), and CCR6+CXCR3-CCR4+ (Th17) cells polarized or sorted from human peripheral blood express the canonical cytokines and transcription factors associated with their corresponding T-helper phenotype (27–31). Th17-lineage cells are also highly plastic and modify chemokine receptor and effector cytokine expression depending on their environment (32, 33). Classical Th17 cells (CCR6+CXCR3-CCR4+) produce primarily IL-17 and are important for neutrophil recruitment and defense at mucosal sites (34); Th1/17 cells (CCR6+CXCR3+CCR4-) share Th1 and Th17 functionality, respond to IL-12 and IL-23, and produce mostly IFNγ (35); Th17 double positive (DP) cells (CCR6+CXCR3+CCR4+) produce IFNγ and low levels of IL-17 (29), and double negative (DN) cells (CCR6+CXCR3-CCR4-) preferentially produce IL-17 and express transcriptional markers of early Th17 development (IL-17F, STAT3), lymph-node homing (CCR7, CD62L), follicular help (CXCR5, BCL6, ASCL2), and self-renewal (LEFI, MYC, TERC) (29, 36).

To enhance the depth of our analyses, we incorporated 13 surface markers that relate to T cell function into a machine learning model for predicting clinical outcomes. The panel included surface markers associated with: activation/proliferation (CD71, HLA-DR, CD38, CD25); degranulation/cytotoxicity (CD94, CD107a); differentiation/memory (loss of CD27, CD28, CD127); Treg/checkpoint inhibitor/exhaustion (CTLA4, PD-1); Tfh/germinal center homing (CXCR5); and mucosal migration (CCR5). While frequencies of T cell subsets could be analyzed individually, the incorporation of expression levels of T cell functional features resulted in over 400 subset-marker pairs and required use of computational methods to analyze this high-dimensional dataset. Using a random forest machine learning model, we determined whether T cell subset frequencies and marker expression levels could distinguish clinical outcomes. When accurate prediction was achieved, feature importance analysis by Gini score (37) determined the most discriminatory features.

Increased frequencies of CD4 TEM Th2 and Th17-lineage subsets and reduced expression of T cell activation/differentiation markers among CT+ or CT+CoP participants compared to uninfected were most informative to the model’s correct classification of CT infection. The model was unable to accurately predict endometrial infection but was successful in identifying features associated with absence of reinfection. The most discriminating features were increased frequencies of CD4 TCM Th1, Th17, Th1/17, and Th17 DN cells and increased expression of markers associated with central memory and Th17-lineage. Although these data suggest a protective role for Th17 cells, their contribution to human immunopathogenesis remains undetermined.

## RESULTS

### CT genital tract infection elicits monocytes, plasma cells, and CD4 T effector memory cells in the periphery

This study acquired CyTOF data from PBMCs of 82 of the 246 participants enrolled in TRAC (21). The sociodemographic characteristics, self-reported STI diagnoses, and exposure histories of this sub-cohort resemble those previously described for the full cohort (Table S1, S2) (21). Study participants were assigned to infection outcome groups according to the presence of CT or copathogens (CoP) *Neisseria gonorrhoeae* (NG) and/or *Mycoplasma genitalium* (MG) at enrollment: Uninfected (CT- NG- MG-), CT+ (CT+ NG- MG-), or CT+CoP (CT+; NG+ and/or MG+). PBMCs, collected at enrollment and 1 month post treatment (1M), were immunophenotyped by CyTOF using a 33-marker panel (Fig 1A, Table S3). The manual gating hierarchy and definitions used to determine frequencies of each cell type are outlined in Table S4. Frequencies of total myeloid cells, B cells, CD4 T cells, and CD8 T cells were similar across all groups (Fig 1B-E), as were dendritic cells and NK cells (data not shown). However, classical, intermediate, and non-classical monocytes were increased in CT+ and CT+CoP women at enrollment and 1M, with significant increases noted in non-classical monocytes, which are important for antigen presentation (Fig 1F-H). Macrophage frequencies were variable but slightly increased in CT+ participants at enrollment and 1M (Fig S1A). Although frequencies of total B cells were not different at enrollment and 1M, frequencies of plasma cells were increased in CT+ but not CT+CoP participants (Fig 1I). Previous studies have shown that CT drives a strong and specific antibody response (21, 38), consistent with a high frequency of plasma cells. Although frequencies of total T cells were not different, frequencies of CD4 TEM cells were significantly increased in CT+ and CT+CoP participants at enrollment and 1M (Fig 1J), while CD8 TEM were only slightly increased in CT+ and CT+CoP participants (Fig 1K). In contrast, CD4 and CD8 TCM frequencies were similar or slightly reduced in CT+ and CT+CoP participants compared to uninfected (Fig 1L, M). Frequencies of CD4 and CD8 stem cell-like memory T cells (TSCM) were moderately decreased in CT+ and CT+CoP participants at enrollment and 1M, likely a reflection of the proportional increase in TEM (Fig S1B, C). Overall, these data demonstrate that CT infection drives increased frequencies of peripheral non-classical monocytes, plasma cells, and CD4 TEM cells, and that responses remain elevated for at least one month after treatment.

**Figure 1.**
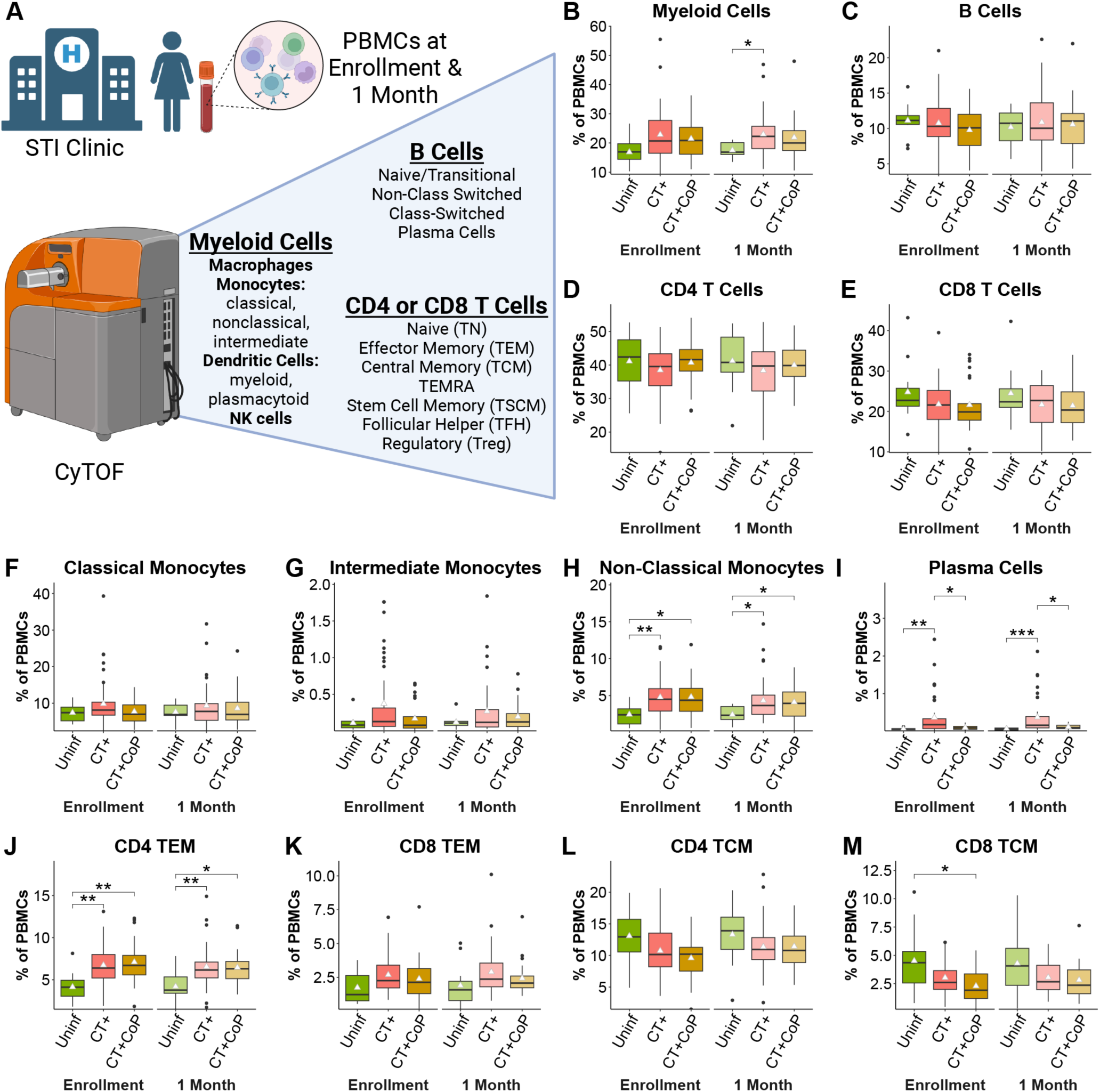
CT genital tract infection elicits monocytes, plasma cells, and CD4 T effector memory cells. (A) Major immune populations defined by manual gating in peripheral blood of 82 participants at enrollment and 1 month post-treatment. (Created with Biorender.com) (B-M) Population percentages of total PBMCs. White triangle represents mean. Significance determined by Dunn’s test with Bonferroni correction for multiple comparisons.

### CT+ participants have increased peripheral frequencies of Th2 and Th17-lineage CD4 TEM cells

We used differential expression of the chemokine receptors CCR6, CXCR3, and CCR4 to define Th1, Th2, and Th17 subsets (Fig 2A, Table S4) (27–29). CD4 T cells, and specifically IFNγ- producing Th1 cells, are broadly accepted as the primary contributors to protection against CT in animals and humans (22, 39), and yet, frequencies of CD4 TEM Th1 cells were not different between CT-infected and uninfected participants (Fig 2B). Instead, frequencies of CD4 TEM Th2 cells were significantly increased in CT+ and CT+CoP participants at enrollment and 1M (Fig 2C), consistent with strong antibody responses detected in response to CT (21, 38). Frequencies of CD4 TEM Th17 and Th17 DN cells were also increased in CT+ participants at enrollment and 1M, whereas there was no difference in frequencies of Th1/17 or Th17 DP cells (Fig 2D-G). These differences were maintained in CT+CoP participants for CD4 TEM Th2 but were dampened in the case of Th17 lineage cells (Fig 2C-F), suggesting that coinfection dilutes Th17 responses elicited by CT.

**Figure 2.**
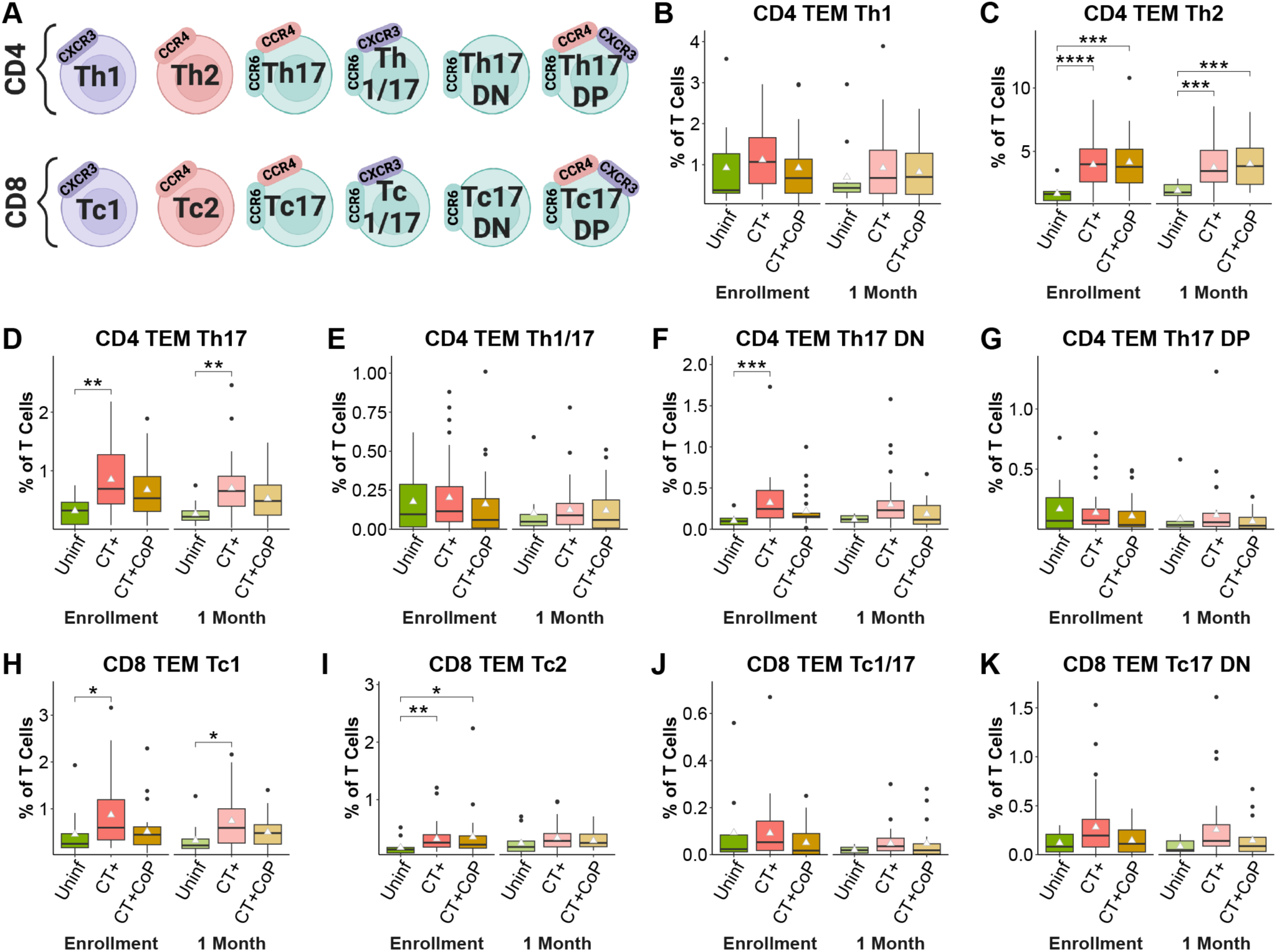
Increased frequencies of Th2 and Th17-lineage CD4 TEM cells among CT+ participants compared to uninfected participants. (A) Subsets of CD4 and CD8 TEM were identified by expression of chemokine receptors (Created with Biorender.com). (B-K) Population percentages of total T Cells. Triangle represents mean. Significance determined by Dunn’s test with Bonferroni correction for multiple comparisons.

We similarly investigated cells comprising the CD8 memory compartment (Tc1, Tc2, Tc17, etc.; Fig 2A, Table S4). Frequencies of CD8 TEM Tc1 and Tc2 were significantly increased in CT+ participants compared to uninfected (Fig 2H-I). There was no change in CD8 TEM Tc1/17 (Fig 2J), but Tc17 DN cells trended toward an increase in CT+ participants (Fig 2K), both of which parallel the results for CD4 TEM subsets. Again, these differences were reduced when comparing uninfected to CT+CoP participants. CD8 TEM Tc17 and Tc17 DP were not detected because there was very low co-expression of CCR6 and CCR4 on CD8 T cells. CD4 or CD8 TCM subset frequencies were similar or slightly decreased in CT+ participants (data not shown), following the trend previously observed for frequencies of total CD4 and CD8 TCM (Fig 1L-M). Overall, these results suggest that CT infection drives a peripheral adaptive immune response dominated by CD4 and CD8 TEM Th2/Tc2 and CD4 TEM Th17-lineage cells.

We then implemented an unsupervised clustering-based strategy to increase the resolution of our findings obtained by supervised manual gating (Fig 3A-F). Clustering with the *k-* means algorithm partitioned cells based on expression of six phenotypic markers (CD45RA, CCR7, CD95, CCR6, CXCR3, CCR4). We previously defined TSCM cells as CD45RA+CCR7+CD28+CD95+ by manual gating (Table S4), but since expression of CD28 was similar across all clusters, this marker was excluded, and TSCM were defined as CD45RA+CCR7+CD95+. Following clustering, memory and T-helper subtypes were annotated based on marker expression as previously (Table S4, S5; Fig 3A-F). Annotation revealed similar clusters to manual gating overall, but enhanced resolution allowed detection of heterogeneity in TN subsets (Fig 3C, F). CD4 clusters 1, 20, and 22 and CD8 clusters 6, 15, and 22 expressed high levels of both CD45RA and CCR7, representing the classical definition of TN. However, CD4 clusters 10, 14, 16 and 19 expressed lower levels of CD45RA and/or CCR7, suggesting that these cells were differentiating from a naïve state to a memory state. Additionally, CD8 clusters 1, 19, 4, and 13 were CD45RA+CCR7+ like traditional TN cells but also expressed CXCR3, a marker typically reserved for memory Th1 cells. Studies have shown that naïve human CD8 T cells expressing CXCR3 have greater potential for effector differentiation (40).

**Figure 3.**
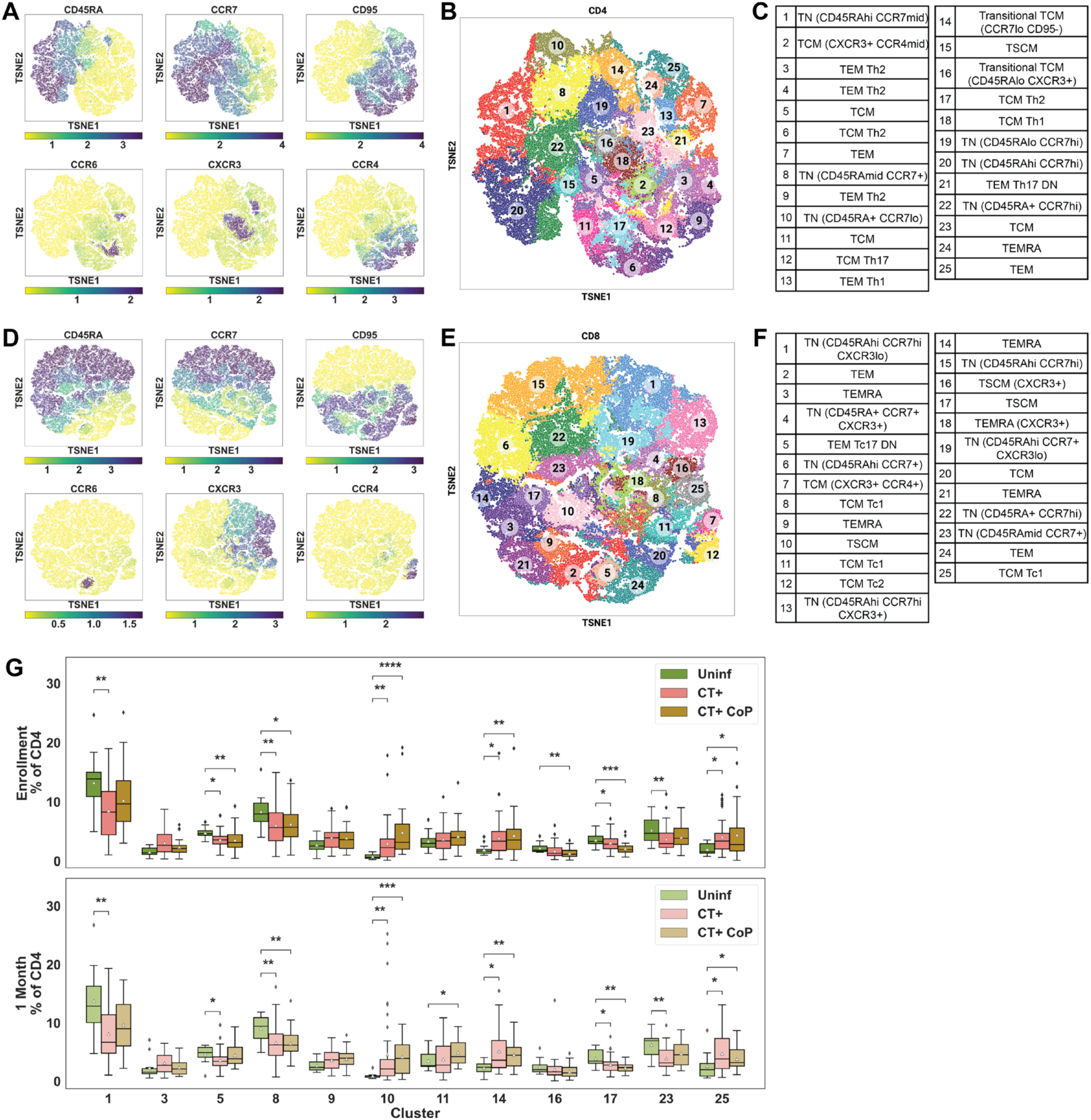
CD4 but not CD8 automated gating frequency features differentiate CT+ and CT+CoP participants from uninfected participants. (A-F) Automated gating by *k*-means clustering and cluster annotation. (A-B) Expression of phenotypic markers (CD45RA, CCR7, CD95, CCR6, CXCR3, and CCR4) was used to partition CD4 T cells into 25 *k*-means clusters. (C) Resulting CD4 clusters were annotated based on marker expression (See Table S4, S11). (D-E) CD8 T cells were partitioned into 25 *k*-means clusters based on the expression of the same phenotypic markers as for CD4 T cells and (F) were annotated. (G) Boxplots representing frequencies of CD4 clusters with significant differences in frequency (Wilcoxon P<0.05). Statistical comparisons between CT+ or CT+CoP and Uninf by Wilcoxon test. White triangle represents mean.

Clusters with frequencies that were statistically significant (P<0.05 by Wilcoxon test) between Uninfected and CT+ or CT+CoP participants at Enrollment or 1 Month are represented in Fig 3G as a distribution of cluster frequencies per participant. Frequencies of CD4 clusters 10 (TN with low CCR7), 14 (Transitional TCM), 25 (TEM), and 3 and 9 (TEM Th2) were increased in CT+ and CT+CoP participants compared to uninfected, while frequencies of clusters 1 and 8 (classical TN), 5 and 23 (TCM), and 17 (TCM Th2) were decreased, suggesting that CD4 T cells respond to CT by evolving away from a naïve state. Similar immune signatures were observed at enrollment and 1 month, suggesting that changes in immune responses in the periphery are maintained after treatment. These findings were consistent with manual gating where overall CD4 TEM (Fig 1J), and especially TEM Th2 cells (Fig 2C), were increased in CT-infected participants compared to uninfected while overall frequencies of CD4 TCM were trending towards decrease (Fig 1L).

### CT+ and CT+CoP participants express reduced T cell activation and differentiation markers on several CD4 and CD8 subsets compared to uninfected participants

While the unsupervised clustering approach lent confidence to our previous findings and uncovered previously unappreciated nuance in the TN population, it demonstrated limited resolution of low frequency T cell subsets defined by manual gating, such as Th17-lineage cells, that were of particular interest to us. Therefore, for further analysis of marker expression on each T cell subset, we used manually gated subsets. We implemented a random forest machine learning model trained by T cell subset frequencies as well as expression levels of the phenotypic markers used above to define T cell subsets (CD45RA, CCR7, CD95, CCR6, CXCR3, CCR4) and 13 surface markers that relate to T cell function to predict clinical outcomes. The panel included surface markers associated with the following T cell functions: activation/proliferation (CD71, HLA-DR, CD38, CD25); degranulation/cytotoxicity (CD94, CD107a); differentiation/memory (loss of CD27, CD28, CD127); Treg/checkpoint inhibitor/exhaustion (CTLA4, PD-1); Tfh/germinal center homing (CXCR5); and mucosal migration (CCR5). The upper quartile expression of each of these 19 markers was calculated for each CD4 or CD8 manually gated subset (as previously defined, Fig 2, Table S4) and for each participant. These upper quartile expression levels (functional features) were then used in addition to the manually gated frequency features to train the random forest model. This machine learning approach allowed interrogation of over 400 features defined by subset-marker pairs.

Classification of uninfected versus CT+ or CT+CoP participants subset frequency and functional features was successful for CD4 (Fig 4A; AUC range 0.96-1.00) or CD8 T cells (Fig4H; AUC range 0.93-0.98). Together, these results suggest that frequency and functional features of CD4 and CD8 T cells contribute to distinction between CT-infected participants and uninfected participants. The similarity between model success at enrollment and 1 month suggests that these signatures were maintained after infection resolution.

**Figure 4.**
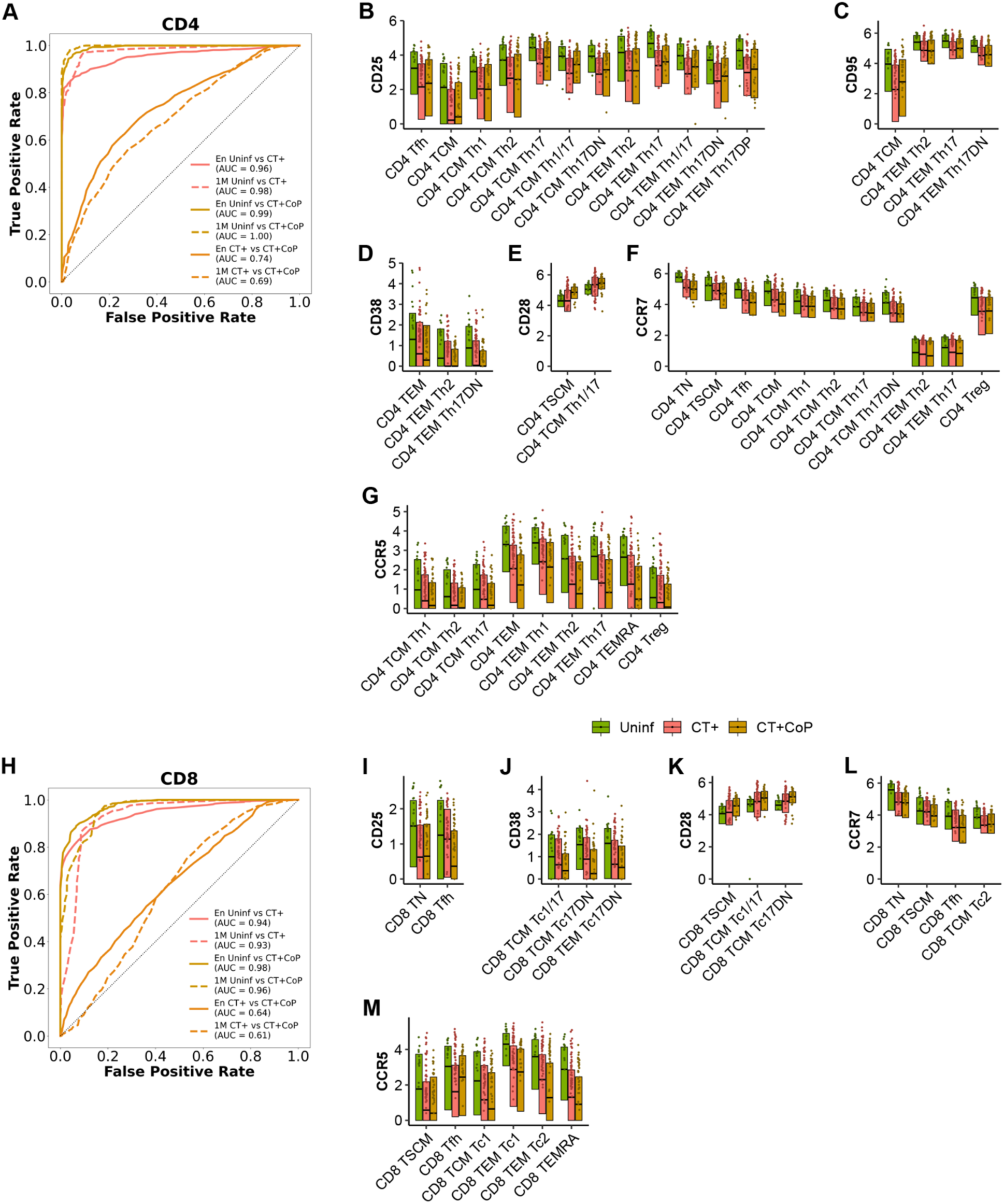
CT+ and CT+CoP participants express reduced T cell activation and differentiation markers on several CD4 and CD8 subsets compared to uninfected participants. A random forest model trained on (A) CD4 or (H) CD8 manually gated subset frequency features and upper quartile expression features was used to discriminate between uninfected, CT+, and CT+CoP outcomes at enrollment (En) or 1 month (1M). Receiver operating characteristic (ROC) curves describe performance of the model. Top markers of (B-G) CD4 or (I- M) CD8 T cells were identified by feature importance analysis (see Suppl tables 6-7). Features were included on the plot if they were statistically significant (as defined by P<0.001 by Wilcoxon rank sum test between outcomes) for at least one comparison between uninfected versus CT+ and/or uninfected versus CT+CoP. Boxplots represent the overall distribution of marker expression. Individual points represent the upper quartile expression for each participant in the group. (AUC = area under the ROC curve)

To identify prominent cellular subsets that contributed to prediction, a measure of feature importance, or Gini score, was assigned to each frequency feature or functional feature (37). Briefly, the Gini score encodes the extent to which each frequency feature or functional feature provides the model with useful information, such that subsets with high Gini scores are likely to be the most useful in distinguishing between clinical outcomes. The Gini scores for predicting classification of infection status are represented in Table S6 for CD4 features and Table S7 for CD8 features. In general, features contributing to accurate classification of uninfected versus CT+ and uninfected versus CT+CoP were similar for the models informed by CD4 and CD8 features, consistent with similar accuracy between the two comparisons and weak classification accuracy for CT+ versus CT+CoP. A subset of upper quartile functional features with highest importance by Gini score and significant difference in expression between uninfected and CT+ or CT+CoP by Wilcoxon test (P<0.001) is shown in Fig 4B-G (CD4), I-M (CD8). Several subsets of both CD4 and CD8 T cells in CT infected participants expressed lower levels of CD25 (IL-2Rα), CD95 (Fas), and CD38, markers associated with T cell activation and antigen experience, than the same subsets in uninfected participants (Fig 4B-D,I-J). Loss of costimulatory molecules CD27 and CD28 is associated with T cell differentiation, and CD28 expression was higher in some CD4 and CD8 subsets from CT+ and CT+CoP participants than in uninfected participants (Fig 4E, K), suggesting that peripheral blood from infected participants contained less differentiated T cells. Activated, antigen experienced, differentiated T cells may have trafficked to the infected genital tract, resulting in a reciprocal decrease in differentiation markers in the blood. Several subsets, especially TSCM and TCM subsets, also expressed lower levels of CCR7 in CT+ or CT+CoP participants than uninfected participants (Fig 4F, L). This aligns with increased frequencies of effector memory (CCR7-) T cells in CT+ and CT+CoP participants and suggests that the central memory cells of infected participants are shifted more toward an effector phenotype. Additionally, expression of the mucosal homing marker CCR5, a marker known to be important for trafficking of T cells to the genital tract during murine chlamydial infection (41, 42), was decreased in several subsets in CT+ or CT+CoP participants compared to uninfected participants (Fig 4G, M). This CCR5 expression pattern could suggest that CCR5+ T cells had migrated to the genital mucosa at the time of sampling. Frequencies of T cell subsets, especially TEM Th2, TEM Th17DN, and TEM Th17 among CD4 T cells (Table S6) and TEM Tc2 and generic TEM among CD8 T cells (Table S7), were also important contributors to the differentiation of CT+ from uninfected participants. Trends in feature importance analysis were similar for both CD4 and CD8 T cells at 1 month (Fig 4, Table S6,7), suggesting maintenance of the T cell signature following infection resolution.

Discrimination of CT+ versus CT+CoP participants using CD4 or CD8 frequency demonstrated only weak to moderate success, with greater success from CD4 features (Fig 4A; AUC range: 0.69-0.74) than CD8 (Fig 4H; AUC range: 0.61-0.64). The top contributing feature by Gini score was higher CTLA4 expression on CD4 T cell subsets among CT+CoP participants compared to CT+ participants (Table S6). CTLA4 (cytotoxic T lymphocyte antigen 4) plays an important role in CD4 Treg function by downregulating T cell activation. NG evades host immunity by inducing TGFβ, which is important for Treg development. The increase in CTLA4 expression on CD4 T cell subsets among CT+CoP participants compared to CT+ participants could be reflective of NG-mediated immune evasion.

### Neither T cell subset frequencies nor functional expression features predict endometrial CT infection

Endometrial biopsies from all participants were tested for CT, NG, and MG infection at enrollment, enabling bisection of the CT+ and CT+CoP groups into CT infections that had ascended to the upper genital tract (Endo+) and CT infections that were limited to the cervix (Endo-). All major cell populations and T cell subsets were defined by manual gating (Table S4), and comparisons of population frequencies performed between Endo+ and Endo- participants. Endo+ participants exhibited significantly increased frequencies of CD4 TSCM, which were not observed in Endo+ CT+CoP participants (Fig S2A). Frequencies of CD8 TEM were not different between Endo+ and Endo- outcomes for CT+ or CT+CoP participants (Fig S2B). Among Endo- participants, there was a trend toward increased frequencies of CD8 TEM Tc17 and TCM Tc17 DN cells (Fig S2C-D), suggesting expansion of these cells may contribute to protection from ascending CT infection. However, the random forest model trained on manually gated subset frequencies with the addition of upper quartile expression of functional markers (Fig S2E-F) failed to accurately classify Endo+ from Endo- participants based on CD4 or CD8 features. Overall, there were relatively minimal differences in the frequency and functional marker expression patterns in peripheral blood T cell subsets between participants with or without ascended CT infection.

### Increased TSCM, CD4 TCM Th1, and CD4 TCM Th17-lineage subsets but decreased CD4 TEM Th2 cells in peripheral blood are associated with absence of CT reinfection

Individuals treated for CT often experience repeat infections (6). Nevertheless, evidence exists for partially protective immunity after infection (23). TRAC study participants were followed for one year to determine if they became reinfected (21). We again bisected CT+ and CT+CoP groups into FU+ (positive for CT by diagnostic test or self-report at any follow-up visit) and FU- (completed at least 3 out of 4 follow-up visits, and CT negative by diagnostic test and self-report at all visits). Five out of 20 participants in the CT+ FU+ group tested positive for CT at 1M, while none of the participants in the CT+CoP FU+ group tested positive for CT at 1M. Removing these 5 participants in the CT+FU+ group from the analysis did not change the overall differences observed between FU- and FU+ participants.

We compared frequencies of manually gated T cell subsets between FU- and FU+ individuals. Frequencies of CD4 and CD8 total T cells and CD4 and CD8 TSCM cells were enriched in FU- participants without co-infection at enrollment and remained expanded 1 month post treatment (Fig 5A-D). CD4 Treg cells were enriched in FU- participants at the enrollment visit (Fig 5E). Interestingly, frequencies of CD4 TEM Th2 cells were increased in FU+ participants (Fig 5F), suggesting that although these cells are induced by CT infection (Fig 2C), they are not protective. In contrast, frequencies of CD4 TCM Th1, Th17, Th1/17, and Th17 DN cells were increased in FU- participants (Fig 5G-J). These data suggest CD4 and CD8 TSCM and CD4 TCM Th1/Th17 lineage cells, are important for natural immunity against incident CT infection. Differences in frequencies of Th17 family subsets between FU- and FU+ participants were only detected among CT+ and not CT+CoP participants, which suggests that Th17 immune responses specific to CT are diluted by coinfection.

**Figure 5.**
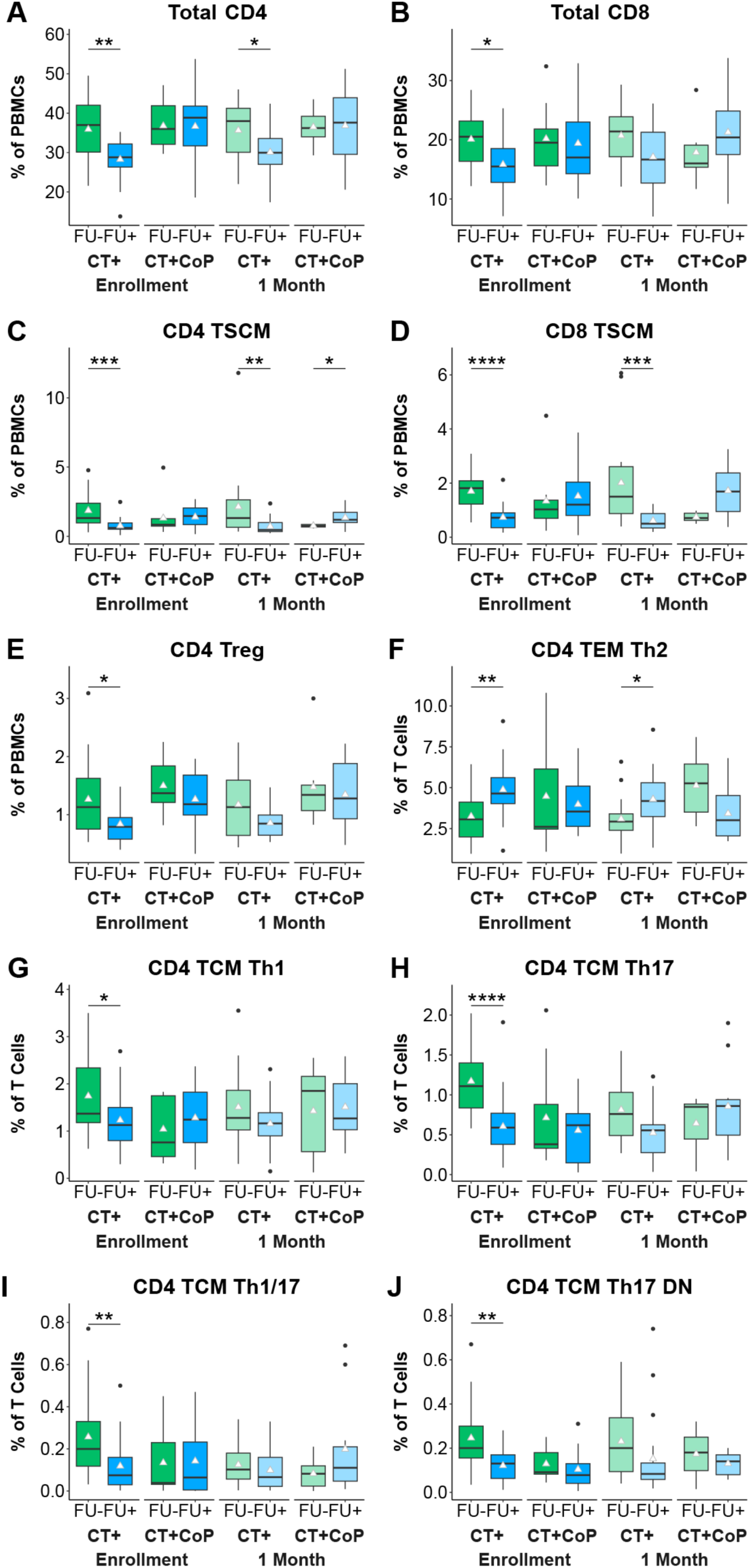
CD4 TCM Th1 and Th17 family subsets are enriched in peripheral blood of participants without documented CT reinfection. Frequencies of manually gated T cell subsets were stratified by follow-up negative (FU-) and follow-up positive (FU+) status. (A-J) T cell subset percentages. White triangle represents mean. Statistical significance determined by Wilcoxon rank- sum test.

We then repeated the comparison using frequencies of clusters generated by *k*-means (Fig 3A-F) and observed trends toward higher frequencies of clusters 15 (TSCM) and 12 and 21 (Th17 family) and lower frequencies of clusters 3, 4, and 9 (Th2 TEM) in FU- participants compared to FU+ (Fig S3; see Fig 3C for cluster annotations). These trends were consistent with results obtained by traditional manual gating (Fig 5) and suggest a protective role for TSCM and TCM Th17-lineage cells but not for Th2 TEM. However, there was limited resolution of Th17 family subsets by clustering compared to manual gating.

### Central memory and Th17-lineage markers distinguish participants who did not become reinfected with CT

We used a random forest machine learning model trained with manually gated T cell subset frequencies and functional expression level features to classify FU+ versus FU- status (Fig 6). Classification of FU+ versus FU- was most accurate for CT+ participants at enrollment (CD4 ROC AUC = 0.72; CD8 ROC AUC = 0.70) (Fig 6A,B). Analysis of feature importances by Gini score identified cell-types and subset-specific expression patterns that were most distinct (Table S8). Among CT+ participants, the most prominent CD4 features contributing to successful FU- versus FU+ classification were decreased frequencies of Th2 cells and increased frequencies of Th17 family subsets in FU- participants (Table S8). The upper quartile functional features with highest importance by Gini score and significant difference in expression between FU+ and FU- by Wilcoxon test (P<0.05) are highlighted in Fig 6C-F. Important CD4 features included increased expression of CCR7 on T cell subsets among FU- participants, consistent with increased central memory, and increased CCR6 expression on Th17 subsets among FU- participants, consistent with increased Th17 lineage (Fig 6C-D). Important CD8 features among FU- participants included increased frequencies of CD8 TSCM, increased expression of CCR7 (similar to the CD4 model), and reduced expression of CD127 (IL-7 receptor, loss of which is associated with differentiation and memory) (Table S8, Fig 6E-F). Overall, the model was most successful in distinguishing FU+ from FU- participants among CT+ participants when informed by CD4 T cell frequencies at enrollment. CD8 T cell features and T cell features at 1 month were less informative, and coinfection with NG or MG diluted the success of the model. Overall, these results suggest a protective role for CD4 TCM Th17-lineage cells against CT reinfection.

**Figure 6.**
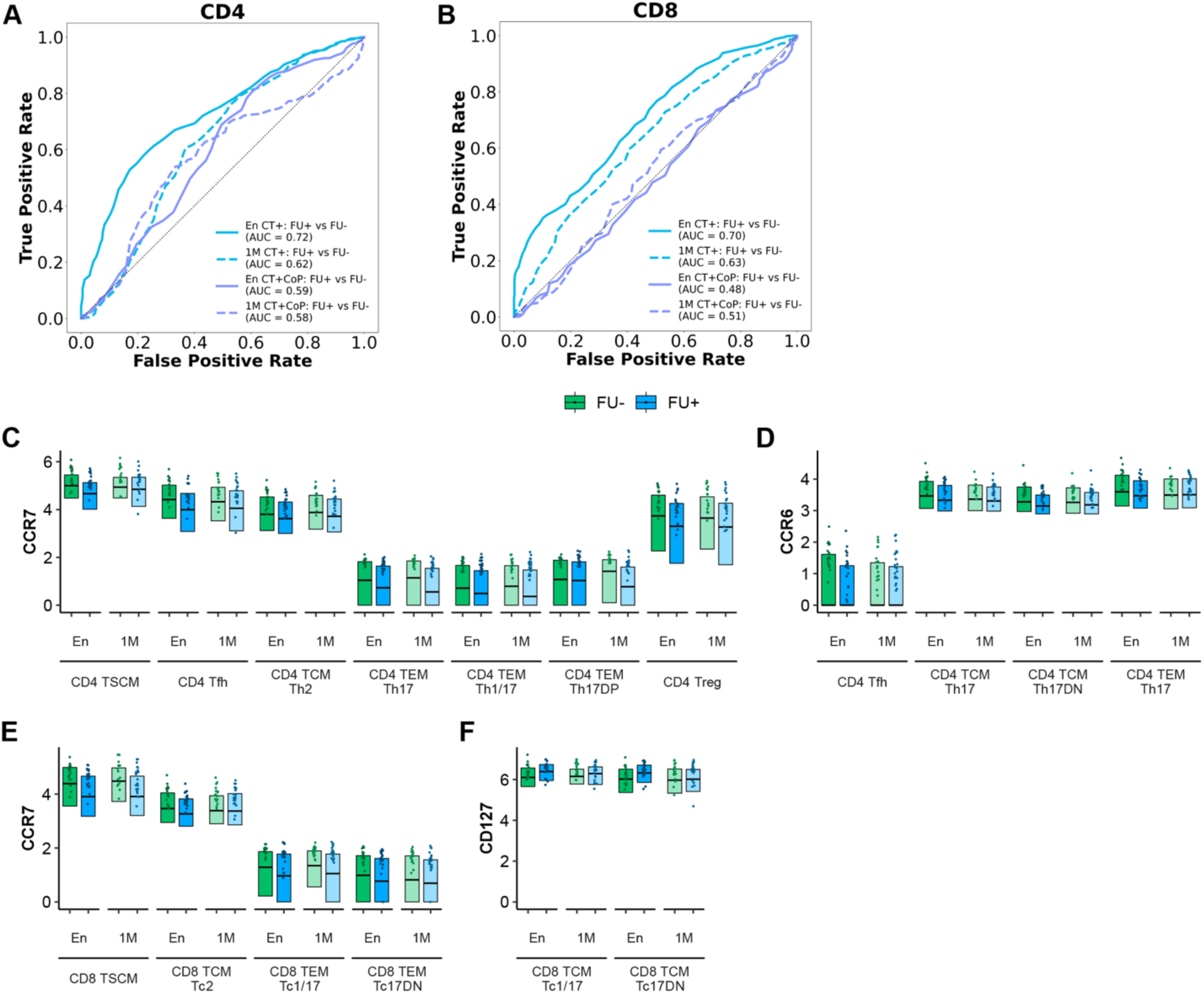
Increased central memory and Th17-lineage markers distinguish participants without documented CT reinfection. A random forest model trained on (A) CD4 or (B) CD8 manually gated subset frequency features and upper quartile expression features was used to discriminate between FU+ and FU- groups at enrollment (En) or 1 month (1M). Receiver operating characteristic (ROC) curves visualize the performance of the model. Top markers of (C-D) CD4 or (E-F) CD8 T cells among CT+ participants were identified by feature importance analysis (see Table S9). Features were included on the plot based on a significance threshold of P<0.05 by Wilcoxon rank sum test for FU+ verus FU- at Enrollment. Boxplots represent the overall distribution of marker expression. Individual points represent the upper quartile expression for each participant in the group. (AUC = area under the ROC curve)

## DISCUSSION

Using samples from a uniquely characterized and clinically relevant cohort, this study immunophenotyped responses to CT infection with high dimensional analysis of surface marker expression on PBMCs and found specific T cell subsets correlated with reduced incidence of CT reinfection. Overall, broad immune cell populations in peripheral blood differed between uninfected and CT+ participants, but were similar between CT monoinfection and coinfection with NG and/or MG. CT+ participants displayed increased frequencies of non-classical (CD14-CD16+) monocytes, important for antigen presentation, T cell proliferation, patrolling tissues for signs of injury, and increased Fc receptor-mediated phagocytosis (43).

While frequencies of immune populations were largely similar in CT+ participants regardless of coinfection, plasma cells were elevated in participants with CT monoinfection, but their frequencies among CT+CoP participants were similar to uninfected. This is consistent with robust antibody responses to CT infection (25, 44, 45), but this response appears suppressed by coinfection with other STIs. NG opacity proteins bind carcinoembryonic antigen-related cellular adhesion molecule 1 (CEACAM1 or CD66a) on B cells, leading to inhibitory signals which promote B cell death (46). Plasma cells also express CD66a (47). Thus, opacity protein interactions with CD66a could contribute to reduced numbers of plasma cells.

CD4 TEM frequencies were strikingly increased in CT+ and CT+CoP participants compared to CD8 TEM frequencies. These data are consistent with our prior investigation of CT antigen specific responses in a subset of TRAC participants where we observed higher levels of IFNγ production and a greater breadth of recognition of CT antigens among CD4 T cells compared to CD8 T cells (22). Lower CD8 TEM frequencies are also consistent with engagement of the PD- 1 receptor on CD8 T cells by antigen presenting cells inhibiting generation of CT-specific memory CD8 T cells (48). Frequencies of TEM but not TCM were increased in CT+ compared to uninfected participants in our study.

Investigation of CD4 TEM subsets revealed that CT+ participants had higher frequencies of peripheral Th2, Th17, and Th17 DN cells compared to uninfected participants, with unchanged frequencies of Th1 cells. CD4 Th1 cells are classically defined by production of IFNγ and expression of CXCR3 and/or CCR5 (28). They are well-documented contributors to chlamydial clearance and protection from reinfection in mouse models (18, 41, 42) and associate with resistance to reinfection in humans (22, 23). We did not find that Th1 cells were expanded in the peripheral blood of CT+ participants, potentially explaining why CT reinfection is common. However, Th1 cells express the chemokine receptors CXCR3 and CCR5, which bind the chemokines CXCL10 and CCL5 (RANTES), respectively (49). We previously reported detecting these chemokines in cervical secretions of TRAC participants (50), and they are highly expressed in genital tract tissues of *C. muridarum* infected mice (41). In this study, we found decreased expression of CCR5 on T cell subsets among CT+ and CT+CoP participants. Thus, it is possible that despite overall expansion of Th1 cells, they were preferentially recruited to the site of infection and consequently not increased in the periphery. In contrast, peripheral CD4 Th2 cells were significantly enriched in CT+ and CT+CoP participants, consistent with high antibody titers to CT infection that fail to prevent reinfection (6, 21, 25).

Th17 cells are highly plastic. They are mainly defined by their expression of CCR6 (51) but can gain or lose chemokine receptors and their associated effector functions (e.g., cytokine production) (32). Th17 cells can be pathogenic, but they are also important contributors to protection against mucosal pathogens (52, 53). We used expression of CCR6, CXCR3, and CCR4 to define four subsets of Th17 family cells: classical Th17, Th1/17, Th17 DP, and Th17 DN (29). We found increased frequencies of classical Th17 and Th17 DN cells in CT+ participants compared to uninfected, which were reduced by coinfection. Frequencies of parallel populations of CD8 TEM cells including Tc2 and Tc17 DN cells were also increased in CT+ and CT+CoP participants, albeit to a lesser extent than CD4 counterparts. Co-expression of CCR6 and CCR4 was extremely rare on CD8 T cells, and therefore Tc17 and Tc17 DP populations were excluded from analysis.

An important goal of the study was to analyze expression of additional surface markers associated with activation/proliferation (CD71, HLA-DR, CD38, CD25); degranulation/cytotoxicity (CD94, CD107a); differentiation/memory (loss of CD27, CD28, CD127); Treg/checkpoint inhibitor/exhaustion (CTLA4, PD-1); Tfh/germinal center homing (CXCR5); and mucosal migration (CCR5) on each T cell subset and to compare expression patterns between clinical outcomes. While T cell subsets could be analyzed individually, investigating differences in expression of phenotypic and functional markers on each subset resulted in over 400 subset- marker pairs. Utilizing a random forest machine learning model allowed us to determine whether T cell subset frequencies and functional expression features were distinct between observed clinical outcomes of CT infection (Uninfected, CT+, CT+CoP), ascension (Endo+/-), or reinfection (FU+/-). If the random forest model was successful using a combination of frequency and functional features to classify clinical outcomes, subsequent feature importance analysis identified which of the T cell subset frequencies and or subset-marker expression pairs were most informative to the model. The model successfully classified CT+ or CT+CoP from uninfected participants using CD4 or CD8 features. Increased frequencies of CD4 TEM Th2, Th17, and Th17 DN cells and distinct expression patterns of CD25, CD95, CD38, CCR7, CD28, and CCR5 were defined as important features for the model. CD25 is the IL-2 receptor and is crucial for T cell activation and proliferation. CD95, or Fas, is involved in apoptosis but is also a marker of antigen experience and stem-cell memory. CD38 is a receptor involved in cell activation, adhesion, and signaling, and is commonly associated with T cell activation. All three of these markers were expressed at lower levels in CT+ or CT+CoP participants than uninfected participants. Lower levels of CCR7 expression on T cells among CT+ and CT+ CoP participants was consistent with higher frequencies of TEM. Additionally, higher levels of CD28 expression on T cell subsets among CT+ or CT+CoP participants suggested less differentiation. Lower levels of CCR5, a molecule important for trafficking of effector T cells to the female genital tract during murine infection (42) and slightly associated with reduced chlamydia burden in humans (54), were also observed on T cell subsets among CT+ or CT+CoP participants. Together, these data suggested that even though CD4 TEM cells were increased in peripheral blood of participants with CT infection, these T cells were not more activated or differentiated than the same subsets in uninfected participants. The chronicity often associated with chlamydial infections could lead to a reduced differentiation state associated with T cell suppression in the periphery, as well as promote ongoing exit of activated, differentiated T cells from the peripheral blood to the infected mucosa. Supporting this hypothesis of reciprocal decrease in activation/homing markers from peripheral blood compared to the site of infection, other studies found that CD38 and/or CCR5 expression was higher among CD4 and CD8 T cells isolated from the genital tract than from peripheral blood of CT+ women (55, 56).

Since CT ascension is a prerequisite for generation of upper tract disease, we sought to identify T cell subsets and expression patterns associated with ascended infection. Peripheral immune cell populations were similar overall between participants with endometrial CT infection and those that had CT infection limited to the cervix. CD8 TEM Tc17 DN cells trended toward increased frequencies in Endo- participants, consistent with our previous study showing that CT- specific CD8 T cells weakly associated with reduced CT ascension (22). Frequencies of CD4 TSCM were significantly increased in Endo+ as well as FU- participants. This is consistent with our prior finding that endometrial infection was associated with reduced risk of follow-up CT infection (21), likely because the increased bacterial burden associated with endometrial infection drives a more robust memory response. However, a random forest model trained with manually gated T cell subset frequencies and functional expression features was unsuccessful in classifying Endo+ from Endo- CT infections, suggesting that immune responses in the periphery are too weak to distinguish whether a participant had ascended CT infection. Evaluation of immune cell populations in the genital tract will likely be necessary to define the specific features of immunity that prevent or limit ascending infection. Future studies will investigate immune cells from cervical and endometrial TRAC samples.

Several T cell subsets were found to be strongly associated with the presence or absence of recurrent CT infection. Sustained increased frequencies of total CD4 T cells in FU- participants may indicate developing natural immunity in some participants. We also detected increased frequencies of CD4 and CD8 TSCM in FU- individuals. TSCM resemble naïve T cells in their expression of CD45RA and CCR7, but they also express CD95, a marker of antigen experience and memory (57). They are minimally differentiated memory cells that have a high propensity for stemness and self-renewal as well as the multipotency to differentiate into a broad range of memory T-helper phenotypes (57). This may explain why they are associated with a FU- outcome. The importance of the nature of the T cell response is demonstrated by the contrasting subsets that were expanded in FU- and FU+ participants. Higher frequencies of TEM Th2 cells, which were strikingly increased in CT+ participants, were associated with a FU+ outcome, while higher frequencies of TCM Th1 cells, which were not different between CT+ and uninfected participants, were associated with FU- status. Overall, an important role for Th1 and Th17-lineage cells in protection from reinfection is suggested by the striking increase in frequencies of TCM Th1 and Th17 cells among FU- participants, along with increased frequencies of TCM Th1/17 and TCM Th17 DN cells. The increased stem-ness and differentiation potential of TSCM and Th17 lineage cells (31, 58), especially Th17 DN (29, 36), is potentially impactful because their plasticity may allow them to adapt quickly to fight infection. A recent study detected enrichment of *Mycobacterium tuberculosis* (Mtb)-specific Th17-like cells in people who were resistant to Mtb infection following exposure compared to those who were latently infected (59). Similar trends were observed when comparing frequencies of T cell subsets as defined by *k*-means clusters. Overall, clusters annotated as TSCM and Th17 lineage were increased in FU- participants and clusters annotated as TEM Th2 were decreased in FU- participants.

Our finding that TCM subsets (TCM Th1, TCM Th17, TCM Th1/17 and TCM Th17 DN cells) rather than TEM subsets were associated with FU- status, is consistent with our detection of CT antigen-specific CD4 TCMs months post treatment among TRAC participants (60). In addition, an analysis of *C. pneumoniae*-specific peripheral blood T cells in healthy seropositive donors found that antigen-specific CD4 TCM were enriched compared to TEM (61). Thus, TCM appear to make up a greater frequency of antigen-specific CD4 T cells important for protection against subsequent infection, although we found that active CT infection drives expansion of TEM cells that remain 1M post-treatment,.

A random forest machine learning model trained to classify FU- versus FU+ informed by manually gated T cell subset frequencies and functional expression features confirmed that the most distinct differences were increased frequencies of TCM Th17, Th17 DN, Th1/17, and Th1 cells and decreased frequencies of TEM Th2 cells in FU- participants. Additionally, we identified distinct expression patterns of surface markers including CCR7 and CCR6. In both the CD4 and CD8 models for CT+ participants, higher expression of CCR7 was observed on several subsets among FU- participants. Since CCR7 is a defining marker of central memory T cells and is important for trafficking to the lymph node, it suggests that central memory cells with the propensity to be long-lived are associated with protection from reinfection. Higher expression of CCR6 on subsets of CD4 T cells among FU- participants suggests an importance not only for Th17 lineage cells, which were defined by the presence of CCR6, but also for its surface expression levels. CCR6 binds CCL20 is highly expressed in mucosal tissues, driving migration of Th17 cells to the site of infection. Additionally, high expression of CCR6 is associated with increased secretion of Th17 effector cytokines and chemokines such as IL-17A, IL-17F, IL-22, and CCL20, and increased expression of *IL17A*, *IL17F*, *IL22*, *IL23R*, *CCL20*, *RORA*, and *RORC* (58).

The finding that CD4 Th1 and Th17 subsets were associated with reduced reinfection informs vaccine development. Different vaccine platforms and adjuvants can shape T-helper responses (62). Vaccination of mice with a recombinant chlamydial antigen with CAF01 (63), or a STING agonist (64) elicited Th1 and Th17 responses that led to reduced chlamydial burden upon challenge. In contrast, a viral vector vaccine failed to elicit antigen-specific CD4 T cell responses or protection (65).

Our study is limited because although we defined immune cell populations increased in CT+ participants compared to uninfected participants, the cells were not restimulated with CT antigens to confirm their CT-specificity. However, immunophenotyping PBMCs directly without restimulation allowed us to confidently define T cell subsets based on chemokine receptors without the risk of downregulation upon restimulation. An important open question is whether Th17 and Th17 DN cells produce CT-specific IL-17 and/or IFNγ, and this should be addressed in future studies. Another limitation of this study is that cell population frequencies could not be correlated with endometrial CT infection. Many of the women participating in TRAC selected contraceptive methods that promote a thin uterine lining, making it difficult to acquire sufficient tissue for evaluation of pathology. However, ongoing studies are investigating local T cell responses in endometrial tissue sections from participants with sufficient tissue.

Overall, we found that frequencies of CD4 TEM, specifically Th2, Th17, and Th17 DN subsets, were enriched in individuals with active CT infection. We also identified that increased frequencies of CD4 TCM Th1, Th1/Th17, and Th17 were associated with individuals who had no documented reinfection over the course of a year. Together, these findings increase our understanding of natural immunity to CT and inform strategies for future vaccine design.

## METHODS

### Sex as a Biological Variable

The TRAC cohort was restricted to persons with a cervix and uterus because the objective of the study was to determine factors associated with ascension of CT from the cervix to the endometrium.

### Study Population

This study involved a subset of 82 of the 246 participants enrolled in the T Cell Response Against Chlamydia (TRAC) cohort (21). TRAC participants were asymptomatic and presented to the clinic for STI screening due to sexual behavior that put them at risk for CT (16). None had signs or symptoms of PID. At enrollment, all participants received antibiotics recommended by the Centers for Disease Control and Prevention for treatment of CT and NG, and individual participants that tested positive for STI at any follow-up visit were retreated (16).

### Definition of Clinical Classifications

#### Uninfected, CT+, and CT+CoP Groups

Nucleic acid amplification testing (NAAT) was used to test cervical swabs collected from study participants at enrollment for CT, NG, and MG. Participants whose swabs tested negatively for all 3 STIs at enrollment were classified as “uninfected”. Participants testing positively for CT but negative for NG and MG were classified as “CT+”. Enrollees testing positively for CT and NG, MG, or both were classified as “CT+CoP”. Of 82 total TRAC participants studied, 12 were uninfected, 44 were CT+, and 26 were CT+CoP (Table S9). Of 26 CT+CoP participants, 11 were CT+ NG+ MG-, 12 were CT+ MG+ NG-, and 3 were CT+ NG+ MG+.

#### Endometrial positive and endometrial negative CT

Endometrial biopsies were also tested by NAAT to determine whether infection was limited to the cervix or had ascended to the upper genital tract. Within CT+ and CT+CoP groups, enrollees testing positively for CT at the cervix but not the endometrial biopsy were classified as Endo-, while those testing positively for CT at both sites were classified as Endo+. No participants tested positive for CT solely in the endometrial sample. Of 44 CT+ participants, 19 were Endo+ and 25 were Endo-. Of 26 CT+CoP participants, 11 were Endo+ and 15 were Endo- (Table S10).

#### Follow-up CT positive and follow-up CT negative

NAAT testing was performed on cervical brush samples at enrollment and at 1, 4, 8, and 12 month follow-up visits. Participants with a positive cervical CT test at enrollment and a positive cervical CT test or self-reported CT diagnosis from an outside clinic in the time since last visit at any follow-up visit (regardless of how many visits they completed) were classified as follow-up positive (FU+). Participants that tested positive for cervical CT at enrollment, completed at least 3 out of 4 follow-up visits, tested negative for cervical CT at all follow-up visits, and self-reported no CT diagnosis from an outside clinic in the time since last visit at any follow-up visit were classified as follow-up negative (FU-). 9 participants attended less than 3 follow-up visits (FU N/A) and were excluded from follow-up analysis. Of CT+ participants, 19 were FU-, 21 were FU+, and 4 were FU N/A. Of CT+CoP pariticpants, 9 were FU-, 12 were FU+, and 5 were FU N/A (Table S10). All uninfected samples were FU- and the group was therefore not bisected.

### PBMC CyTOF

Peripheral blood mononuclear cells (PBMCs) were isolated from whole blood via a density gradient using lymphocyte separation media (Corning). Cells were frozen in 90% FBS with 10% DMSO and stored in liquid nitrogen until thawed for analysis. Dead cells were stained with 1 μM Cell-ID Cisplatin 198-Pt (Fluidigm). Cells were stained with an antibody cocktail comprising 33 antibodies (Table S3) and human Fc block (eBioscience) in cell staining buffer (CSB, Fluidigm), followed by staining with secondary antibody (Anti FITC-144Nd). Stained cells were fixed with 2% PFA in PBS followed by 1:2000 Cell-ID Intercalator-Ir-125 μM (Fluidigm) in fix and perm buffer (Fluidigm). Cell concentrations were adjusted to 0.5 x 10^6^ cells/mL in CAS (Cell Acquisition Solution; Fluidigm) with Four Element Calibration beads (Fluidigm). Samples were acquired on a Helios instrument (Fluidigm) by the UNC Mass Cytometry Core.

### Data Analysis

#### Defining T cell subsets by manual or automated gating

Raw CyTOF files were cleaned using the Pathsetter algorithm (66).

##### Manual gating

Cleaned FCS files were analyzed in Cytobank (Beckman Coulter) and FlowJo (BD) software. The manual gating strategy and surface marker designations selected for immune cell phenotypes are summarized in Table S4. Some gated CD8 populations were undetectable (<0.1% of T cells) and were excluded from further analysis (Table S4).

##### Automated gating

Batch effects were removed from the dataset using the combat algorithm (67). A random subset of 15,000 cells was selected from each sample, resulting in 1,230,000 total cells for further analysis. We then performed automated cell-population discovery followed by feature engineering, as outlined in Stanley et al. (68). CD4 and CD8 T cell populations were each partitioned into 25 clusters using *k*-means clustering. Briefly, each cell was represented by expression of six phenotypic markers (CD45RA, CCR7, CD95, CCR6, CXCR3, CCR4), which were input to the clustering algorithm. Moreover, each cell was labelled in terms of one of the 25 clusters, based on its expression across these six phenotypic markers (Table S5). Modeled after canonical immune features that are often computed based on manual gates, we engineered per-cluster frequency features, where for each sample, we computed the proportion of its cells assigned across each of the 25 clusters.

#### Feature engineering

While previous studies have created functional features based on the mean or median expression of functional markers in each cluster (68, 69), some functional markers in this dataset were expressed by a small proportion of cells, such that there were some cases where median expression of a marker was zero. Therefore, we computed the upper quartile expression of each marker across each manually gated T cell subset for each sample. For samples with 0.1% events assigned to a particular subset, the upper quartile features were imputed with the mean of each subset-marker feature. Additionally, subset-marker features for which upper quartile expression was below 2 (range 0-500) were excluded from the training dataset. Upper quartile features were arcsinh transformed with co-factor 5. In all, 19 frequency features and 233 upper quartile expression features were included in the CD4 model training dataset, and 14 frequency features and 174 upper quartile expression features were included in the CD8 model training dataset.

#### Model training

We trained random forest models using engineered manually gated frequency features and upper quartile expression features to classify and predict clinical outcomes for each participant. For each clinical outcome comparison, we performed 30 trials of five-fold cross validation, whereby each fold used 80% of samples for training and 20% for testing. Reported area under the ROC curves (AUCs) represent the mean accuracy across 30 trials. For biological interpretability of each of the trained random forest models, we further computed per-feature importance measures with the Gini index (37). A higher per-feature Gini score implied that the feature was important in the clinical prediction task. We further imposed directionality to the computed per-feature Gini scores to readily identify condition-specific, predictive features by first choosing a particular clinical outcome to be the *positive (negative) direction*, such that we multiplied the Gini score by 1 (-1), if it was associated with the positive (negative) direction.

### Statistics

Statistical calculations were computed in R and python. Statistical tests used are outlined in the appropriate figure legends. P<0.05 was used to determine a significant result. Significance symbols used in figures are defined as: *P<0.05, **P<0.01, ***P<0.001, ****P<0.0001.

### Study Approval

The institutional review boards for human research of the University of Pittsburgh and the University of North Carolina approved the study protocol. All participants provided written informed consent at the time of enrollment and agreed to be contacted to return for follow-up visits 1, 4, 8, and 12 months after enrollment.

### Data and Code Availability

CyTOF data are available for download via ImmPort at https://www.immport.org under study accession number SDY2772. Manually gated T cell subset frequencies reported in the Figures are available in Supporting Data Values. Code for (1) automated gating, (2) feature engineering, (3) random forest training and Gini feature importance analysis, and (4) upper quartile expression boxplots are available via GitHub at https://github.com/ksyount/TRAC_PBMC_CyTOF.

## Supporting information

Supporting Data Values

## AUTHOR CONTRIBUTIONS

KSY, AK, XZ, NS, and TD designed research studies. KSY and AK conducted experiments. HCW and SLH collected participant samples and performed diagnostic testing. TP advised regarding mass cytometry panel design. KSY, CC, CL, NVM, and CBB analyzed data. KSY, CC, NS, CMO, and TD wrote the manuscript. All authors reviewed the manuscript.

## ACKNOWLEDGEMENTS

We thank the women who agreed to participate in this study; Ingrid Macio, Melinda Petrina, Carol Priest, Abi Jett, and Lorna Rabe, for their efforts in the clinic and the microbiology laboratory; and the staff at the Allegheny County Health Department STD Clinic. We thank Jason Whitmire for critical review of the manuscript. We also thank Marie Iannone and the UNC Mass Cytometry Core, which is supported by the University Cancer Research Fund (UCRF) and UNC Cancer Center Core Support Grant #P30CA016086. This work was supported by the National Institute of Allergy and Infectious Diseases through U19AI144181 (TD) and R01AI170959 (CMO). KSY received funding from UNC STI/HIV postdoctoral training grant T32AI007001 and the American Association of Immunologists Intersect Fellowship Program for Computational Scientists and Immunologists.

