## Supplemental Figures for "Unique T cell signatures associated with reduced *Chlamydia trachomatis* reinfection in a highly exposed cohort"

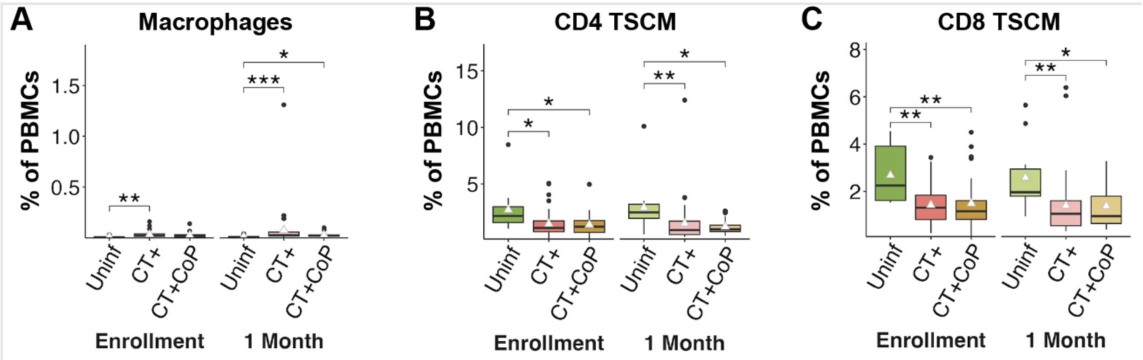


**Supplemental figure 1. Frequencies of manually gated cell subsets among participants.** Population percentages of total PBMCs. White triangle represents mean. Significance determined by Dunn's test with Bonferroni correction for multiple comparisons. (Uninf = CT- and NG- and MG-; CT+ = CT+ and NG- and MG-; CT+CoP [Co-Pathogen] = CT+ and NG+ and/or MG+)

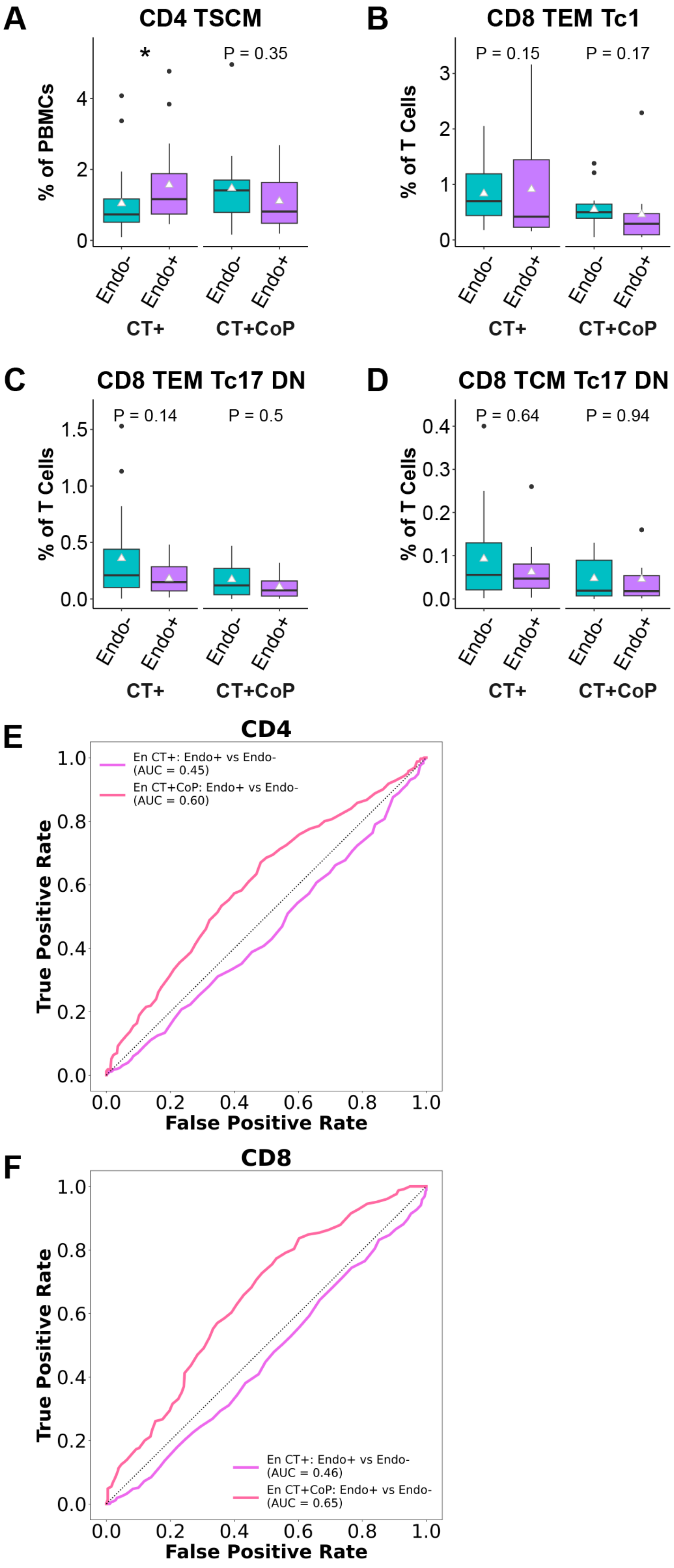


**Supplemental figure 2. Features of CD4 and CD8 T cells are not distinct between CT+ participants with or without infection that ascended to the endometrium.**

Frequencies of manually gated T cell subpopulations were stratified by cervix only CT (Endo-) or cervix and endometrial CT (Endo+) status. (A-D) T cell subset percentages. White triangle represents mean. Significance determined by Wilcoxon rank-sum test. A random forest model trained on (E) CD4 or (F) CD8 manually gated subset frequency features and upper quartile expression features was used to discriminate between Endo+ and Endo- groups at enrollment (En). Receiver operating characteristic (ROC) curves visualize the performance of the model. (AUC = area under the ROC curve).

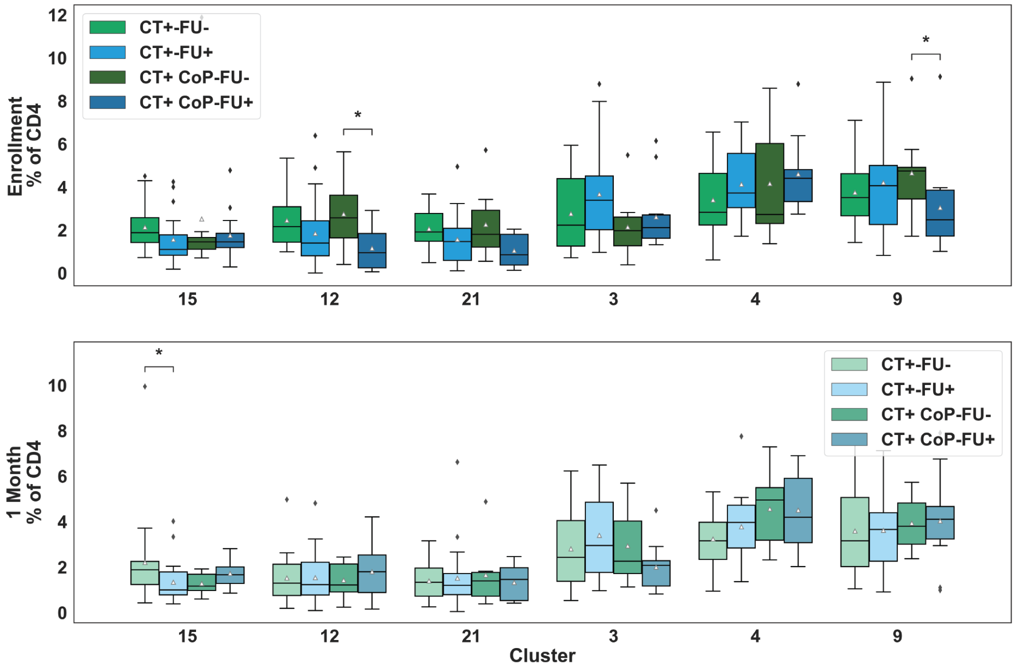


**Supplemental figure 3. Differences in CD4 T cell subset frequencies between FU- and FU+ participants by automated gating are similar to those observed by manual gating.**

Boxplots representing frequencies of select CD4 clusters (reference Fig 3A-C) among participants. Statistical comparisons between FU- and FU+ by Wilcoxon test. White triangle represents mean.
